# Cell wall forming chitin synthases in a chytrid fungus

**DOI:** 10.1101/2025.08.12.668609

**Authors:** Trupti Prakash Gaikwad, Michael Cunliffe

## Abstract

Chitin is a critical structural component of fungal cell walls, yet our understanding of its synthesis across the kingdom Fungi remains limited. Here, we investigate chitin synthase diversity, transcription and localisation in the saprotrophic chytrid *Rhizoclosmatium globosum* (*Rg*), expanding insights into fungal cell wall biology beyond Dikaryan models. We identified 20 chitin synthase genes in the *Rg* genome, including canonical Division I and II types, and a distinctive chitin synthase gene containing a glycoside hydrolase domain linked to β-glucan synthesis. Transcriptomic analysis through zoospore, germling and immature thallus developmental stages revealed stage-specific expression patterns, with active gene diversity correlating with increasing morphological complexity. Using electroporation-based transformation and fluorescent fusion constructs, we demonstrated successful expression and localisation of two chitin synthases during cell development. Localisation patterns showed dynamic redistribution from cytoplasmic dispersion in early encysted cells to concentrated signals at the sporangium wall. Expression in the apophysis and at the apophysis–sporangium junction indicates the importance of these structures in cell maintenance. Our findings highlight functional specialisation among chitin synthases and underscore the importance of cell wall integrity in chytrid development. This work establishes *Rg* as a genetically tractable model for studying chytrid cell biology and contributes to broader understanding of fungal evolution and cell wall dynamics.

## Introduction

Fungi are one of the most globally prevalent and impactful group of organisms (Case et al. 2025). Key to fungal success is a dynamic cell wall that controls viability, morphology and functionality, through a range of biophysiochemical properties including strength and elasticity (Gow and Lenardon 2023). Chitin, a β-1,4-linked homopolymer of *N*- acetylglucosamine, is a major strengthening structural component of most fungal cell walls and chitin synthases are key in fungal cell wall formation, adding *N*-acetylglucosamine monomers to the growing polymer at the plasma membrane (Gow et al. 2017).

Fungal chitin synthase genes fit broadly into three divisions based on domain architecture and phylogenetic analysis (Fernandes et al. 2016). Division I chitin synthases include both type 1 (Pfam01644, CS1) and type 2 (Pfam03142, CS2) chitin synthase domains, along with a chitin synthase N-terminal domain (Pfam08407, CS1N). Division II chitin synthases have CS2 plus additional domains and Division III chitin synthases have only CS2 domains (Li et al. 2016).

Current understanding of the biology of the fungal cell wall is primarily made from studies of Dikaryan fungi (Ascomycota and Basidiomycota), and of which, most are focused on human- and agricultural plant-associated models. The Chytridiomycota (chytrids) are a major group of fungi that are evolutionarily and biologically distinct from Dikaryan fungi (Figure 1A). Found throughout terrestrial, marine, freshwater and artificial habitats, chytrids are important saprotrophs and parasites of other organisms including algae and amphibians (Berbee et al. 2017, Grossart et al. 2019, Medina and Buchler 2020). Despite their prevalence and importance, there remains a limited understanding of chytrid cell biology including their cell walls (Laundon and Cunliffe 2021).

**Figure 1.**
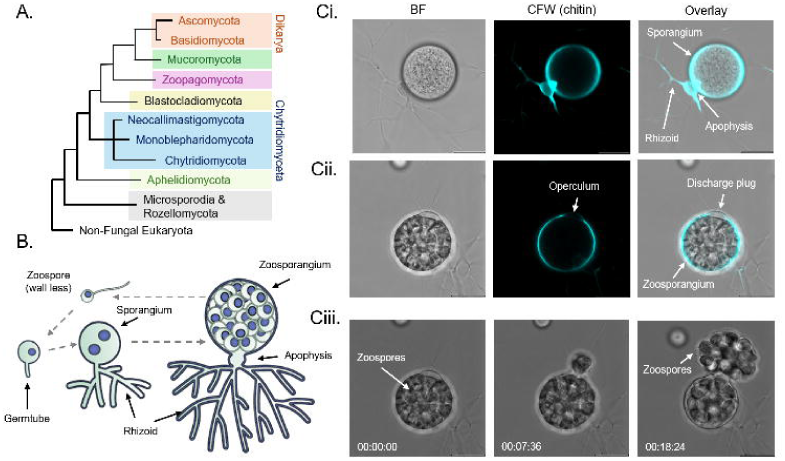
Schematic phylogeny of the kingdom Fungi (based on (Li et al. 2021)) (A). Key stages of the chytrid cell cycle (zoospores > germling > immature thallus > mature zoosporangium) (adapted from (Laundon and Cunliffe 2021)) (B). The chytrid *Rhizoclosmatium globosum* (*Rg*) in brightfield (BF) and confocal microscopy with cell wall chitin labelled with calcofluor-white (CFW) (C). Mature cell showing key features (Ci). Gravid zoosporangium at the point of zoospore release with discharge plug blocking operculum (Cii). Time-series frames from video of zoospore release through the operculum in the cell wall (Supplementary Video 2) (Ciii). Scale bar = 10 µm.

Chytrids are commonly monocentric, developing from a zoospore to a single celled thallus made up of a spherical sporangium for reproduction connected to tubular rhizoids for substrate/host attachment and feeding (Figures 1 B-C, Supplementary Video 1). Within mature sporangia (zoosporangia), wall-less uniflagellate zoospores are produced that are usually motile when released (Figure 1C, Supplementary Videos 1-2). After swimming, and in some species crawling, to find a suitable substrate or host, chytrid zoospores retract the flagellum and encyst by initiating cell wall formation, developing into a walled thallus with an emerging germ tube from which the rhizoid grows (Figure 1B, Supplementary Video 1).

Here, our approach is to use the saprotrophic chytrid *Rhizoclosmatium globosum* (*Rg*) (Figure 1C) to explore chitin synthase genes and domain architecture with transcription through the dynamic cell cycle. Our step change is the development and application of chitin synthase fluorescent fusion constructs successfully transformed into *Rg* to demonstrate the expression and localisation of selected chitin synthases during the chytrid cell cycle.

## Materials and Methods

### Strain and culture conditions

We used the *Rg* strain JEL800 (a model saprotrophic chytrid (Laundon et al. 2020, Laundon et al. 2022, Chrismas et al. 2025)). For all experiments, *Rg* was maintained at 22 °C in the dark on peptidised milk, tryptone and glucose (PmTG) medium (Barr 1986) either as liquid or on agar plates (15 g l^-1^).

### Plasmids and transformation via electroporation

The plasmids pEK36, pEK39 and tdTomato-Lifeact-7 used previously with *Batrachochytrium dendrobatidis* (*Bd*) were obtained from the Addgene plasmid repository. pEK36 and pEK39 were a gift from Lillian Fritz-Laylin (Addgene plasmids 210598 and 210599) (Kalinka et al. 2024) and tdTomato-Lifeact-7 was a gift from Michael Davidson (Addgene plasmid 54528) (Fiolka et al. 2012) (Supplementary Table 1). In this study, we also made new plasmids by isolating genomic DNA from *Rg* with the DNeasy Blood & Tissue Kit (Qiagen) and PCR amplifying the *Rg* ORY39038 promoter with Phusion high-fidelity DNA polymerase (New England Biolabs). We generated *Rg* ORY39038 promoter mClover, mRuby3 and tdTomato expression constructs by seamless cloning (NEBuilder® HiFi DNA Assembly, New England Biolabs, Cat. No. E5520S) with digested pEK36, pEK39 and tdTomato-Lifeact-7 (Supplementary Table 2). All plasmid sequences were verified with PCR amplification using Phusion high-fidelity DNA polymerase (New England Biolabs) and whole plasmid sequencing (Source Bioscience).

For the transformation experiments, 10 ml overnight grown *Escherichia coli* TOP10 in Luria-Bertani medium containing each plasmid was followed by plasmid preparations with a miniprep kit (QIAprep Spin Miniprep Kit) following manufactures instructions. Transformation of *Rg* followed the protocol developed for *Bd* (Swafford et al. 2020, Kalinka et al. 2024) with minor modifications as follows. Plasmids were diluted in 15 µl RNA-free water and mixed with 25 µl electroporation buffer AS27 (25 mM D-Mannitol, 0.7 mM MgCl_2_ and 1mM KCl).

Additional AS27 buffer, 1% tryptone and 1 mm electroporation cuvettes (Geneflow Electroporation Cuvettes, E6-0050) were held on ice. Zoospores were harvested from 24 hr old lawns on agar plates and washed three times in 10 ml chilled AS27 buffer with centrifugation for 5 min at 2,500 g and 4 °C. The washed zoospore pellet was resuspended in 100 µl AS27 buffer, abundance determined with a Luna Cell Counter (Luna Automated Fluorescence Cell Counter, Luna-FL (L20001)) and made to a working abundance of 3 x 10^7^ zoospores ml^-1^. To electroporate, 40 µl of zoospores and 40 µl of plasmid working solution were added to a pre-chilled cuvette, before the cuvette was held on ice for a further 10 min. The cuvette contents were mixed by pipetting 3-4 times and the cuvette placed in the electroporator (Bio-RAD-Gene Pulser XCell). Impedance was measured followed by two positive polarity 350 V/1 ms poring pulses at 100 ms intervals, with an immediately following 20 V/50ms polarity-exchanged transfer pulse. The cuvette was placed on ice for 10 min and then 200 µl chilled 1% tryptone added. Following a further 10 min on ice, 400 µl chilled 1% tryptone added, mixed by pipetting and the contents of the cuvette transferred to a 5 ml tube (Falcon) containing 3.9 ml chilled 1% tryptone before being separated into 600 µl aliquots in 24-well tissue culture plates for recovery. After recovery for 18 hr, dependent on the plasmid transformed (Supplementary Table 2), either hygromycin B (Gibco) or neomycin (Gibco) dissolved in 1% tryptone were added to a final concentration of 20 µg ml^-1^ and 3 mg ml^-1^ respectively and incubated for a further 78 hr.

### Cell fixation, staining and microscopy

Chitin was labelled with calcofluor white (CFW) staining (Laundon et al. 2020). For actin staining with Alexa fluor 488 phalloidin, 200 µl PmTG grown cells were added to poly-l-lysin coated glass-bottom dishes for 15 min before the medium was removed and the attached cells were washed three times with 500 µl phosphate-buffered saline (PBS) and then fixed with 4% formaldehyde for 1 hr. The fixed cells were again washed three times with PBS and then once with 100 mM PEM buffer (100 mM PIPES (piperazine-N,N′-bis(2- ethanesulfonic acid)) buffer at pH 6.9, 1 mM EGTA (ethylene glycol tetraacetic acid), and 0.1 mM MgSO_4_). Cells were then stained with 150 nM Alexa fluor 488 phalloidin (Thermo Fisher) in PEM buffer for 30 min at room temperature in the dark, followed by washing three times with PEM buffer.

Cells were imaged with a Leica SP8 confocal microscope (Leica) with acquisition settings as follows: CFW excitation/emission 405/426-470 nm (intensity 0.098 %, gain 10), mRuby and tdTomato 561/570-620 nm (intensity 2.7 %, gain 651), mClover3 488/500-650 nm (intensity 1 %, gain 113) and Alexa fluor 488 phalloidin 488/450-570 nm (intensity 1.1 %, gain 77). All comparisons between cell stages were made under identical acquisition settings. Alexa fluor 488 phalloidin and tdTomatolifeact7 images are maximum intensity projections at 0.52 µm z-intervals. All imaging was at room temperature. Images were processed using LasX software (Leica).

### Identification and analysis of chitin synthases

To identify putative chitin synthase genes, the *Rg* genome (Mondo et al. 2017) was initially screened for glycosyl transferase family 2 genes through the MycoCosm database (Grigoriev et al. 2013) targeting the conserved type 1 (Pfam01644, CS1) and type 2 (Pfam03142, CS2) chitin synthase domains. All resulting amino acid sequences from reciprocal open reading frames were further assessed for associated Pfam matches via the Conserved Domain Database (Wang et al. 2022). Sequence alignments and phylogenetic analysis followed (Li et al. 2016), using a combination of MUSCLE and the Maximum likelihood method in MEGA (Tamura et al. 2007). Division I chitin synthases were aligned based on the combined CS1N, CS1 and CS2 domains. Division II were aligned based only the CS2 domains.

### Transcription of chitin synthases

RNA sequencing (RNA-Seq) was used to determine the presence and quantity of chitin synthase gene mRNA at zoospore, germling and immature thallus stages of the *Rg* cell cycle. The RNA-Seq data used are from a previous study characterising the general *Rg* cell cycle (Laundon et al. 2022). In summary, synchronised *Rg* cultures (n = 3) were grown with PmTG medium and harvested at 0 hr (zoospore), 1.5 hr (germling), 10 hr (immature thalli). Cell stages were confirmed with microscopy. RNA was extracted using the RNeasy extraction kit (Qiagen). Sequencing was carried out using Illumina NovaSeq 6000 technology and base calling by CASAVA by Novogene (https://www.novogene.com). Raw reads were filtered for adaptor contamination and low-quality reads before being mapped against the *Rg* JEL800 genome using HISAT2 and used to calculate Fragments Per Kilobase per Million mapped fragments (FPKM).

### Chitin synthase constructs and transformants

Plasmids for expression and localisation of chitin synthase (ORY52914 and ORY53285) fluorescent fusion proteins were made using seamless cloning (NEBuilder HiFi DNA Assembly, New England Biolabs) following the same protocol as above (Supplementary Table 2).

## Results

### Demonstration of Rg transformation and fluorescent protein expression

Plasmids previously used in the transformation of *Bd* (Kalinka et al. 2024) with the non- native *Spizellomyces puncatus* (*Sp*) histone 2B promoter (Medina et al. 2020) demonstrated cytosolic mClover (pEK36) and mRuby3 (pEK39) fluorescent protein expression in *Rg* under hygromycin B selection (Figure 2A). We also successfully transformed and showed expression of the tdTomato-Lifeact-7 fusion plasmid in *Rg* under neomycin selection (Figure 2B). LifeAct is a 17-amino-acid peptide that stains filamentous actin (F-actin) (Riedl et al. 2008). Co-localisation of actin-specific fluorescent phalloidin with tdTomato fluorescence in *Rg* confirmed LifeAct fidelity (Figure 2B). As well as demonstrating transformation and expression of fluorescent proteins in *Rg* with plasmids previously used for *Bd* and *Sp* with non-*Rg* promoters, we also made three modified constructs with a *Rg* promoter. Using the *Rg* promoter from the putative chitin synthase ORY39038, we demonstrated cytosolic mClover (ORY39038mClover3) and mRuby3 (ORY39038mRuby3) fluorescent protein expression under hygromycin B selection and similarly tdTomato (ORY39038 tdTomato) fluorescence under neomycin selection (Figure 2C).

**Figure 2.**
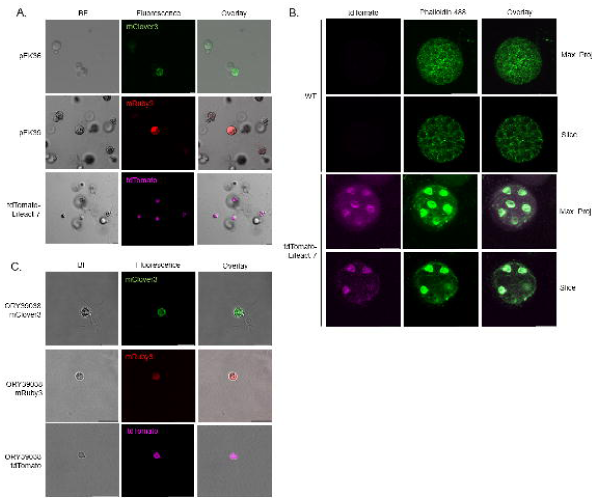
Successful transformation of *Rhizoclosmatium globosum* (*Rg*) with functional fluorescent proteins mClover (pEK36), mRuby3 (pEK39) and tdTomato (tdTomato-Lifeact-7) (A). tdTomato-LifeAct localisation showing the distribution polymerised actin in *Rg* zoosporangium. Representative maximum intensity projections and single slice images of wild type (WT) and tdTomato-LifeAct (purple) fixed and stained with phalloidin 488 (green) (B). Successful transformation of functional fluorescent proteins in *Rg* promoter control (ORY39038) (C). Brightfield (BF). Scale bar = 10 µm.

### Chitin synthase genes and domains

Twenty chitin synthase genes were predicted in the *Rg* genome based on the presence of the conserved CS1 and CS2 domains. Of which, 11 are likely members of Division I because of the characteristic CS1N, CS1 and CS2 domain structure and no other associated domains (Figure 3A). The other genes lacked CS1 domains and formed three groups (Figure 3B).

**Figure 3.**
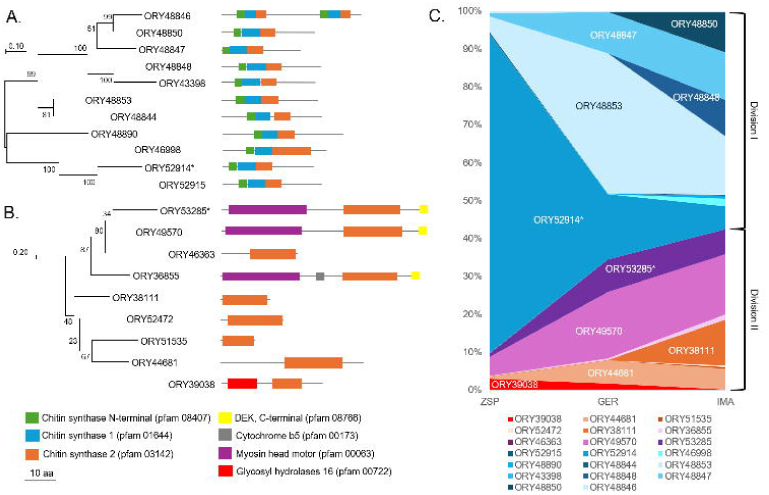
Maximum likelihood phylogenies of *Rhizoclosmatium globosum* (*Rg*) Division I (A) and Division II (B) chitin synthases. Relative proportions (% of FPKM) of chitin synthase gene mRNA transcripts between the *Rg* zoospore (ZSP), germling (GER) and immature thallus (IMA) cell stages (Absolute FPKM in Supplementary Table 3). Asterisks show chitin synthases ORY52914 (CS1N, CS1 and CS2) and ORY53285 (MMD, CS2 and DEKC) used for fusion constructs.

One group made up of 4 genes, contained CS2 domains and additional domains with 3 including N-terminal myosin motor domains (Pf00063, MMD) and DEK C-terminal domains (Pf08766, DEKC), and 1 also having a cytochrome b5-like domain (Pf00173, Cyt-b5). One of the 4 genes clustered in the group had no additional domains (i.e. CS2 only). A second group made up of 4 genes other than CS2 had no additional domains. ORY39038 is distinctive compared to the other *Rg* chitin synthases, containing both a CS2 domain that branched separately from the other CS2 domains and a glycoside hydrolase 16 domain (Pf00722, GH16).

### Transcription of chitin synthases through cell development

The abundance (FPKM) and relative proportions (% of FPKM) of chitin synthase gene mRNA transcripts varied between the *Rg* cell stages, with increasing diversity of transcribed genes matching growing cell morphological complexity (Figure 3C, Supplementary Table 3). In zoospores, Division I ORY52914 (CS1N, CS1, CS2) dominated the mRNA transcripts, then becoming less abundant in the other cell stages. The change from the wall-less zoospores to walled germlings corresponded with an increase in the range of major chitin synthase mRNA transcripts from other Division I and Division II genes. The immature thalli (i.e. developed rhizoid with no obvious zoosporegenesis) had the greatest range of chitin synthase gene mRNA transcripts compared to the less morphologically complex germlings and zoospores.

### Expression and localisation of chitin synthases

To determine the expression and localisation of the Division I ORY52914 (CS1N, CS1 and CS2) and Division II ORY53285 (MMD, CS2 and DEKC), fusion constructs under native promoter control were generated (ORY52914-mRuby3 and ORY53285-mRuby3) and transformed (Figure 4A and 4B). In both transformants, mRuby3 signal was distributed throughout the cytoplasm of the early encysted cell. As the cell developed to the germling stage (i.e. smaller sporangium with initial rhizoid development), both the ORY52914-mRuby3 and ORY53285-mRuby3 mRuby3 signals were concentrated along the out edge of the sporangium wall and to some extent the rhizoid with ORY53285-mRuby3, corresponding with the CFW signal. During the germling stage, increased ORY52914-mRuby3 mRuby3 signal was also associated with the chitin-rich area at the sporangium-germtube/rhizoid junction (Figure 4A). In the immature thallus (i.e. sporangium expanding in size but with no obvious zoospores), the mRuby3 signals in both ORY52914-mRuby3 and ORY53285- mRuby3 remained along the sporangium edge matching the CFW defined cell wall. Also at this stage, the ORY52914-mRuby3 mRuby3 signal was concentrated in the apophysis (Figure 4A). At the immature thallus stage, the ORY53285-mRuby3 mRuby3 signal was instead associated with the chitin-rich pseudo septum at sporangium and apophysis junction (Figure 4B). In later stage of the cell cycle, when the zoosporangium (i.e. zoospores visible) stopped increasing in size, the ORY53285-mRuby3 signal associated with the sporangium wall was relatively reduced compared to the previous stage whilst the distinctive mRuby3 signal associated with the sporangium and apophysis pseudo septum remained (Figure 4B).

**Figure 4.**
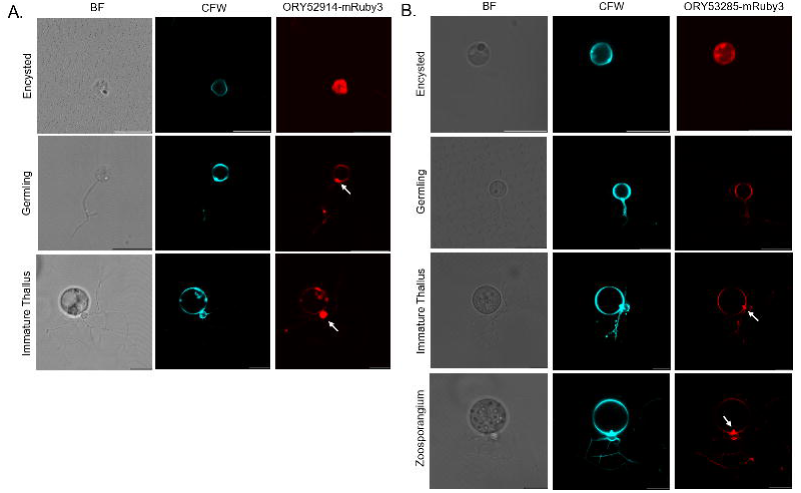
*Rhizoclosmatium globosum* (*Rg*) expressing constructs ORY52914-mRuby3 (Division I CS1N, CS1 and CS2) (A) and ORY53285-mRuby3 (Division II ORY53285 MMD, CS2 and DEKC) (B) observed with brightfield (BF) and confocal microscopy with the cell wall chitin labelled with calcofluor-white (CFW). Arrows highlight in the germling stage the sporangium-germtube/rhizoid junction with increased ORY52914-mRuby3 signal, the immature thallus apophysis also with increased ORY52914-mRuby3 mRuby3 signal and the chitin-rich pseudo septum at sporangium and apophysis junction with increased ORY53285-mRuby3 signal in the immature thallus and zoosporangium stages. Scale bar = 10 µm.

## Discussion

Perhaps the major finding from our study is the demonstration of the expression and localisation of chitin synthases in a chytrid cell, with distribution changing through the walled cell development cycle. For both fusion constructs, mRuby3 signal was dissipated in the cytoplasm of the early stages of the chitin-containing walled encysted cell. In the following germling and subsequent stages, the chitin synthase mRuby3 signals became evenly distributed at the outer wall edge, in accordance with the archetypical plasma membrane association of fungal chitin synthases and corresponding with the observed isotropic growth of the expanding sporangium part of the *Rg* cell (Supplementary Video 1) (Laundon et al. 2022).

As well as chitin synthase at the sporangium wall, we also show that within the *Rg* apophysis has chitin synthase present, implying a role of the apophysis in cell wall maintenance of the chytrid cell. Despite being common across the Chytridiomycota, the apophysis has not been well studied and there is a limited understanding of the biological role of the structure in the chytrid cell (Laundon and Cunliffe 2021, Laundon et al. 2022). Through a combination of 3D model reconstruction with serial block-face scanning electron microscopy and live-cell imaging of FM1-43 endomembrane, we have previously shown that the apophysis is enriched with endomembrane and vacuoles and is an active site for intracellular trafficking between the rhizoids and the sporangium via an annular pore (discussed further below). As part of the intracellular trafficking function, the apophysis could have a role in mediating chitin synthases for cell wall formation that warrants focused study.

We also show that the chitin-rich wall region circling the annular pore connecting the apophysis and sporangium has localised chitin synthase expression. This pore and associated chitin-rich wall is the link between the substrate attaching and feeding rhizoid and the sporangium for reproduction (Laundon et al. 2022). The Division II ORY53285 includes a myosin motor domain (Pf00063, MMD) (Figure 3B) which have been studied in Dikaryan fungi (Fernandes et al. 2016). In the corn smut *Ustilago maydis* (Basidiomycota), chitin synthase associated MMD interacts with the actin components of hyphae supporting polar growth (Schuster et al. 2016) and has a role in tethering chitin synthases in position in the cell (Schuster et al. 2012). Actin cables run through the *Rg* rhizoid (Laundon et al. 2020) and in the closely related *Chytriomyces hyalinus* (Chytriomycetaceae) have been shown to pass through the rhizoids and base of the sporangium from the septum (Dee et al. 2019). It is possible that the localisation of the MMD-containing chitin synthase (ORY53285) at this position in the chytrid cell is facilitated by actin. Given that chitin adds mechanical strength to the fungal wall (Gow 2025), our observation of prolonged localised chitin synthase expression and chitin-enrichment at this part of the chytrid cell wall suggests that maintenance of the strength of this junction is important for the cell including during the critical zoosporegenesis stage. Future work could consider the mechanical forces on the chytrid cell. We propose that the apophysis-sporangium junction is a potential ‘weak point’ overcome by reinforcement with chitin.

Chitin synthase gene copy number is variable across the fungal Kingdom (Gonçalves et al. 2016, Li et al. 2016, Liu et al. 2017) and the biological reasoning for the variation is not yet fully understood (Gow 2025). For example, the fission yeast *Schizosaccharomyces pombe* (Ascomycota) has only one chitin synthase gene (Gonçalves et al. 2016) whilst the rice blast *Magnaporthe oryzae* (Ascomycota) has seven chitin synthase genes (Kong et al. 2012). Surveys of chitin synthase genes across the fungal kingdom have shown that non-Dikaryan fungi including chytrids have higher numbers of chitin synthase genes compared to Dikaryan fungi ((Gonçalves et al. 2016, Li et al. 2016, Liu et al. 2017), as we show here with *Rg*.

Most of the *Rg* chitin synthase genes fit within the canonical view of fungal chitin synthase gene domain structures, such as the likely Division I genes made up of the conserved CS1N, CS1 and CS2 domains found widely across the kingdom (Liu et al. 2017). The additional domains accompanying some of the CS2 domains have also been found in representatives across the kingdom (Li et al. 2016). Three of the CS2 domain containing chitin synthase genes had N-terminal myosin motor domains (MMD), which, as discussed above, in Dikaryan fungi have been linked to chitin synthase interaction with cytoskeleton actin and tethering the enzyme to specific locations in the cell (Steinberg 2011, Schuster et al. 2012, Schuster et al. 2016). With the DEK C-terminal domains (DEKC) and cytochrome b5-like domain (Cyt-b5) we show here in *Rg*, though commonly found with Division II chitin synthases, specific roles in chitin-related wall formation in fungi are currently unknown (Li et al. 2016).

Contrasting with the largely canonical chitin synthase genes, one of the *Rg* chitin synthase genes was made up of a distinctive CS2 domain and associated GH16 domain (ORY39038). We have previously suggested that this gene could be involved with *Rg* cell wall β-glucan production, an observation supported by measurements of β-glucan in *Rg* biomass and the fungal β (1,3) glucan synthesis inhibitor caspofungin causing deformed rhizoid growth (Laundon et al. 2020). In Dikaryan fungi, β (1,3) glucan is the other major cell wall polysaccharide and alongside the strength from chitin controls the elasticity of the cell wall matrix (Gow 2025). Along with further work on chitin in chytrid cell walls, other core structural polysaccharides (e.g. β glucans) and potential wall ‘decorating’ polysaccharides need attention.

Transcription of chitin synthase genes changed through *Rg* cell development with a positive relationship between the diversity of transcribed chitin synthase genes and increases in cell morphological complexity (i.e. zoospore>germling>immature thallus). In the multicellular *M. oryzae*, transcription of the seven chitin synthases varies between different major cell types (e.g. vegetative hyphae, appressoria) and when infecting rice leaves indicating that specific chitin synthases have different cellular roles (Kong et al. 2012). For example, specific chitin synthases have been linked to roles in the *M. oryzae* rice infection process, such as the CHS7 gene function in appressorium formation (Kong et al. 2012).

Half of the chitin synthase genes in the *Rg* genome were not detected or minimally detected (≤ 5 FPKM) in the transcriptomes of any of the three stages (Supplementary Table 3). It is possible that some of these genes are active during the mature zoosporangium stage when zoosporegenesis is taking place including having a role in remodelling the cell wall of the zoosporangium to accommodate the discharge plug for zoospore release (Figure 1C, Supplementary Video 2). Chitin synthase gene duplication, including within species duplication, is particularly common in non-Dikaryan fungi including chytrids (Li et al. 2016) and is it therefore likely some degree of functional redundancy is taking place as we indicate here.

The kingdom Fungi covers a vast diversity of the tree of life with a rich evolutionary history in which the Chytridiomycota are a large and important group. Here we expand the toolbox for studying chytrids by applying electroporation-based genetic transformation and the development of expressing and localising fluorescent protein fusions. Along with the freshwater saprotroph *Rg* (this study), the soil saprotroph *Sp* (Medina et al. 2020) and the amphibian pathogen *Bd* (Kalinka et al. 2024), together this creates opportunity for expanding our understanding of the biology, ecology and evolution of chytrids, the wider fungal kingdom and general eukaryote cell functioning.

## Data availability

All PCR primers and plasmids produced in this study are described in Supplementary Table 2. Chitin synthase gene transcriptome data (FPKM) used for Figure 3 (% FPKM) are provided in Supplementary Table 3. All raw transcriptome reads were deposited in the Sequence Read Archive (PRJNA789147) from a previous study (Laundon et al. 2022).

## Acknowledgements

We thank Joyce E. Longcore (University of Maine) for providing *Rg* JEL800, now held at the Collection of Zoosporic Eufungi at the University of Michigan (CZEUM). We thank Glen Wheeler (Marine Biological Association) for helpful advice with confocal microscopy.

Supplementary video 1 was produced by Davis Laundon as part of his PhD studies at the Marine Biological Association for which we thank.

## Author contributions

T.P.G and M.C. conceptualised the study. T.P.G and M.C. acquired and analysed the data. T.P.G and M.C. wrote original drafts, reviewed and edited subsequent drafts. Funding was acquired by M.C.

